# Comparative analysis of gene prediction tools for viral genome annotation

**DOI:** 10.1101/2021.12.11.472104

**Authors:** Enrique González-Tortuero, Revathy Krishnamurthi, Heather E. Allison, Ian B. Goodhead, Chloë E. James

## Abstract

The number of newly available viral genomes and metagenomes has increased exponentially since the development of high throughput sequencing platforms and genome analysis tools. Bioinformatic annotation pipelines are largely based on open reading frame (ORF) calling software, which identifies genes independently of the sequence taxonomical background. Although ORF-calling programs provide a rapid genome annotation, they can misidentify ORFs and start codons; errors that might be perpetuated and propagated over time. This study evaluated the performance of multiple ORF-calling programs for viral genome annotation against the complete RefSeq viral database. Programs outputs varied when considering the viral nucleic acid type versus the viral host. According to the number of ORFs, Prodigal and Metaprodigal were the most accurate programs for DNA viruses, while FragGeneScan and Prodigal generated the most accurate outputs for RNA viruses. Similarly, Prodigal outperformed the benchmark for viruses infecting prokaryotes, and GLIMMER and GeneMarkS produced the most accurate annotations for viruses infecting eukaryotes. When the coordinates of the ORFs were considered, Prodigal scored high for all scenarios except for RNA viruses, where GeneMarkS generated the most reliable results. Overall, the quality of the coordinates predicted for RNA viruses was poorer than for DNA viruses, suggesting the need for improved ORF-calling programs to deal with RNA viruses. Moreover, none of the ORF-calling programs reached 90% accuracy for annotation of DNA viruses. Any automatic annotation can still be improved by manual curation, especially when the presence of ORFs is validated with wet-lab experiments. However, our evaluation of the current ORF-calling programs is expected to be useful for the improvement of viral genome annotation pipelines and highlights the need for more expression data to improve the rigor of reference genomes.

## Introduction

The field of viromics—the characterization of viral communities and populations (viromes) in a given environmental niche (1)—is rapidly evolving along with the increasing discovery and characterization of new viruses across all domains of life (2,3). The development of sequencing technologies, the associated reduction in costs and increased throughput, has made high quality viral metagenomic studies possible (4). As a result, the number of new sequenced virus and phage genomes is expanding at an impressive rate (5,6), arguably without the concomitant improvement in appropriate bioinformatic tools required to examine viral contigs and genomes (7) or to address the number of viral sequences that share little or no homology to any genes of predictable function (uncultivated virus genomes) (6).

New viruses are usually annotated using de novo genome annotation pipelines such as RAST (8), Prokka (9), VIGA (10), and Cenote-Taker 2 (11). All these bioinformatic tools strongly rely on open reading frame (ORF) calling software, such as GLIMMER (12), the GeneMark family of programs (13-15) and Prodigal (16), which are the most commonly used programs. These ORF-calling programs identify genes and their start codons without considering the taxonomical background of the sequence. Although most of these programs were designed for bacterial genome analysis, they have also been used to rapidly annotate complete viral genomes. However, this approach can produce poorly optimized results. For instance non-coding ORFs might be misidentified as coding ORFs, real ORFs might be missed, or start codons misidentified (17). This problem is particularly relevant as the annotation of new viruses relies on previous annotations of similar viruses, resulting in the perpetuation and propagation of annotation errors over time (5).

Recent benchmarking exercises have evaluated the performance of multiple ORF-calling programs for temperate bacteriophage annotation (5,18). However, these investigations relied solely on the genomes of temperate phages whose genes were known empirically. Salisbury and Tsourkas (2019) only considered sequences of *Escherichia* virus Lambda and *Mycobacterium* virus Patience (5), whereas Lazeroff *et al* (2021) performed benchmarking using a total of eight virus genomes, including the aforementioned Lambda and Patience (18); yet, the sample size was smaller than the estimated sample size required for the complete collection of sequenced viral genomes. In fact, when considering all complete bacteriophage genome sequences present in the NCBI Reference Sequence Database (RefSeq), (4,166 at the time of writing) the estimated minimum sample size was 352 (95% confidence interval; 5% error margin) or 3,331 (99% confidence interval; 1% error margin). Similarly, for all complete virus genome sequences reported in RefSeq (13,778 at the time of writing), the estimated minimum sample size was 374 (95% confidence interval; 5% margin of error) or 7,538 (99% confidence interval; 1% error margin) (19).

This study evaluates the performance of multiple ORF-calling programs for viral genome annotation using the whole RefSeq viral database (20). To assess the impact of ORF misidentification, several factors were considered: A false ORF might be treated as a coding sequence, a true ORF might be lost during the bioinformatic prediction process, or the location of start codons was incorrect during the ORF prediction process. The number of ORFs and their coordinates were considered. Unfortunately, despite their importance in viral biology, this benchmarking exercise was not able to include non-coding RNA elements, which have only recently been annotated in viruses (21,22). Rigorous and regular evaluation of such systems in this way is fundamental to the evolution of viral genome annotation pipelines that can keep pace with the ever-increasing volume of virus sequence data.

## Material and methods

### Benchmark creation: database and ORF-calling programs

The RefSeq viral database (20) was used as a gold standard to evaluate the performance of the different ORF calling programs,. The RefSeq collection provides a curated, non-redundant, stable database for annotated reference genomes of viruses, microbes, organelles, and eukaryotic organisms (23). At the time of writing, RefSeq contained 13,778 sequences, of which only 8,267 sequences were complete genomes, 9,505 belonged to viruses infecting eukaryotic host cells, 4,166 belonged to bacteriophages (including 10 sequences of Mollicutes bacteriophages) and 107 were identified as viruses infecting archaeal host cells.

All 13,778 viral genome sequences from RefSeq were submitted to Prodigal v. 2.6.3 (16), GLIMMER v. 3.02 (12), GeneMarkS v. 4.32 (14), PHANOTATE v. 1.5.0 (24), Metaprodigal v. 2.6.3 (25), FragGeneScan v. 1.31 (26), MetaGeneAnnotator (MGA) (27), and AUGUSTUS v. 3.4.0 (28). Prodigal, GLIMMER and GeneMarkS are the most commonly used ORF-calling programs for prokaryotic genomes, being the most critical step for the majority of the de novo bioinformatics pipelines (8,9,29). PHANOTATE was included because it was specifically designed for bacteriophage genome annotation (24). Metaprodigal, FragGeneScan and MGA are particularly useful for metagenomics and metaviromics datasets as they have been optimized for gene identification in highly fragmented assemblies (especially for contigs less than 20,000 bp long) (25-27). All programs were run using the same parameters, focusing especially on the use of the NCBI genetic code 11 (“Bacterial, Archaeal and Plant Plastid Code”) for archaeal viruses and non-Mollicutes bacteriophages, 4 (“Mold, Protozoan, and Coelenterate Mitochondrial Code and Mycoplasma/Spiroplasma Code”) for Mollicutes phages, and 1 (“Standard Genetic Code”) for eukaryotic viruses. In the case of AUGUSTUS, the in-built models for *Staphylococcus aureus, Escherichia coli* and *Homo sapiens* and default parameters were considered for ORF calling. All program outputs were processed using customized Python 3 scripts to retrieve the number of genes and the coordinates of these ORFs.

### Statistical analyses

To evaluate each ORF-calling program, two different analyses were performed: i) coding sequence number prediction, and ii) coding sequence coordinate prediction. First, linear models were used to infer the accuracy or trueness, defined as the proximity of the retrieved number of viral ORFs from every program to the expected number of viral ORFs according to those described in RefSeq for the same virus. Linear models also considered the precision (measurement of the deviation between the retrieved number of viral ORFs for every program and the expected value from the linear model) of the ORF-calling programs in determining the number of viral coding sequences compared to the reference database. All linear models were forced to have intercept zero. The slope was used as a measure of accuracy, while the coefficient of determination (R^2^) was used to measure the precision. Secondly, the prediction quality of the coordinates of the viral coding sequences was evaluated by the F1 score or Sørensen-Dice coefficient, where the precision and sensitivity was defined as:

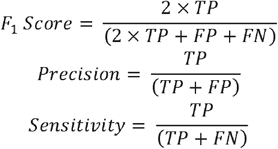

TP indicates the number of true positives (ORFs for which coordinates where exactly the same in both the output file and the reference), FP the number of false positives (ORFs for which coordinates appeared only in the output file) and FN the number of false negatives (ORFs for which coordinates appeared in the reference and were missed in the output file) (17). False Discovery Rate (FDR) and False Negative Rate (FNR) were considered to measure the type I (false coordinates were considered as true coordinates) and the type II (true coordinates were considered as false coordinates) errors. All statistical analyses were performed in R. v. 4.1.0 (30).

### Data availability/Novel Programs, Software, Algorithms

All Python 3 and R scripts are freely available under the GNU General Public License v. 3.0 at https://github.com/EGTortuero/Benchmarking_ORF_calling_programs_in_viral_genomes

## Results

The outputs from each annotation program were evaluated according to two different parameters: (1) number of coding sequences and, (2) coordinates of coding sequences.

### Coding Sequence Number Prediction

Firstly, the accuracy and the precision of the number of viral coding sequences were estimated using linear models. Accuracy was measured by the slope, and precision was measured according to the R^2^ of the regression model. In a general overview, the programs delivered different estimates of the number of coding sequences (Table 1). PHANOTATE, Prodigal and Metaprodigal overestimated the number of ORFs by 30.69%, 1.59% and 1.00% respectively, while the remaining programs tended to underestimate the number of ORFs—the median percentage of underestimation was 26.95 % ± 28.95 %. Despite such observation, Prodigal and Metaprodigal showed the most accurate predictions, being closest to the ideal accuracy of 100.00% (Fig. 1A). However, MGA, Prodigal and FragGeneScan were the three most precise programs according to their coefficients of determination (96.32%, 95.65% and 95.59%, respectively; Table 1). When compared according to host domain, similar results were found for all scenarios tested (Figs. 1B-D). Prodigal outperformed the accuracy test for viruses infecting archaea and bacteria (96.61% and 99.56%, respectively), while GLIMMER and GeneMarkS were the most accurate ORF callers for viruses infecting eukaryotes (99.36% and 97.82%, respectively; Table 1). Finally, when considering the viral nucleic acid, all programs predicted differences in the number of coding sequences (Figs. 1E-F). In fact, while for double-stranded (ds-) and single-stranded (ss-) DNA viruses the most accurate programs were Prodigal (101.59%) and Metaprodigal (101.01%); FragGeneScan (99.62% accuracy and 88.06% precision) and Prodigal (99.01% accuracy and 87.65% precision) generated the most accurate and precise results for ds- and ss-RNA viruses (Table 1).

**Table 1.** Accuracy and precision in the number of coding sequences

| Case (number of sequences) | Program | Accuracy (Slope) | Precision ($R^2$) |
| --- | --- | --- | --- |
| All viruses (13,778) | PRODIGAL | 1.01587 | 0.9565 |
|  | METAPRODIGAL | 1.01004 | 0.9506 |
|  | GLIMMER | 0.70951 | 0.7423 |
|  | GeneMarkS | 0.73051 | 0.7315 |
|  | PHANOTATE | 1.30686 | 0.855 |
|  | MGA | 0.9571 | 0.9632 |
|  | FragGeneScan | 0.95786 | 0.9559 |
|  | AUGUSTUS (*S. aureus*) | 0.83911 | 0.8969 |
|  | AUGUSTUS (*E. coli*) | 0.61472 | 0.8377 |
|  | AUGUSTUS (*H. sapiens*) | 0.43244 | 0.7892 |
| Viruses infecting archaeal hosts (107) | PRODIGAL | 0.96607 | 0.997 |
|  | METAPRODIGAL | 0.9337 | 0.995 |
|  | GLIMMER | 0.50471 | 0.9138 |
|  | GeneMarkS | 0.71465 | 0.7085 |
|  | PHANOTATE | 1.15295 | 0.9899 |
|  | MGA | 0.9133 | 0.9949 |
|  | FragGeneScan | 0.86562 | 0.9933 |
|  | AUGUSTUS (*S. aureus*) | 0.61412 | 0.8888 |
|  | AUGUSTUS (*E. coli*) | 0.4999 | 0.7508 |
|  | AUGUSTUS (*H. sapiens*) | 0.36808 | 0.7855 |
| Bacteriophages (4,166) | PRODIGAL | 0.99555 | 0.9897 |
|  | METAPRODIGAL | 0.98786 | 0.9895 |
|  | GLIMMER | 0.5837 | 0.9386 |
|  | GeneMarkS | 0.61961 | 0.6906 |
|  | PHANOTATE | 1.13126 | 0.9862 |
|  | MGA | 0.98438 | 0.9894 |
|  | FragGeneScan | 0.93567 | 0.9877 |
|  | AUGUSTUS (*S. aureus*) | 0.8262 | 0.9112 |
|  | AUGUSTUS (*E. coli*) | 0.68659 | 0.9232 |
|  | AUGUSTUS (*H. sapiens*) | 0.39223 | 0.8675 |
| Viruses infecting eukaryotic hosts (9,505) | PRODIGAL | 1.06201 | 0.8993 |
|  | METAPRODIGAL | 1.06079 | 0.8857 |
|  | GLIMMER | 0.99358 | 0.7113 |
|  | GeneMarkS | 0.97815 | 0.8566 |
|  | PHANOTATE | 1.70524 | 0.8109 |
|  | MGA | 0.897 | 0.9041 |
|  | FragGeneScan | 1.0089 | 0.903 |
|  | AUGUSTUS (*S. aureus*) | 0.87165 | 0.8721 |
|  | AUGUSTUS (*E. coli*) | 0.45585 | 0.6494 |
|  | AUGUSTUS (*H. sapiens*) | 0.52323 | 0.7378 |
| **ds- and ss-DNA viruses (7,564)** | PRODIGAL | 1.01588 | 0.9565 |
|  | METAPRODIGAL | 1.01007 | 0.9507 |
|  | GLIMMER | 0.70949 | 0.7422 |
|  | GeneMarkS | 0.73032 | 0.7315 |
|  | PHANOTATE | 1.30676 | 0.8551 |
|  | MGA | 0.95717 | 0.9633 |
|  | FragGeneScan | 0.95784 | 0.9559 |
|  | AUGUSTUS (*S. aureus*) | 0.83915 | 0.897 |
|  | AUGUSTUS (*E. coli*) | 0.61482 | 0.8378 |
|  | AUGUSTUS (*H. sapiens*) | 0.43242 | 0.7892 |
| **ds- and ss-RNA viruses (6,214)** | PRODIGAL | 0.99008 | 0.8765 |
|  | METAPRODIGAL | 0.96462 | 0.8703 |
|  | GLIMMER | 0.74766 | 0.8785 |
|  | GeneMarkS | 1.05858 | 0.8735 |
|  | PHANOTATE | 1.4917 | 0.7337 |
|  | MGA | 0.84354 | 0.811 |
|  | FragGeneScan | 0.99624 | 0.8806 |
|  | AUGUSTUS (*S. aureus*) | 0.76804 | 0.804 |
|  | AUGUSTUS (*E. coli*) | 0.3849 | 0.6401 |
|  | AUGUSTUS (*H. sapiens*) | 0.46998 | 0.7512 |

**Figure 1.**
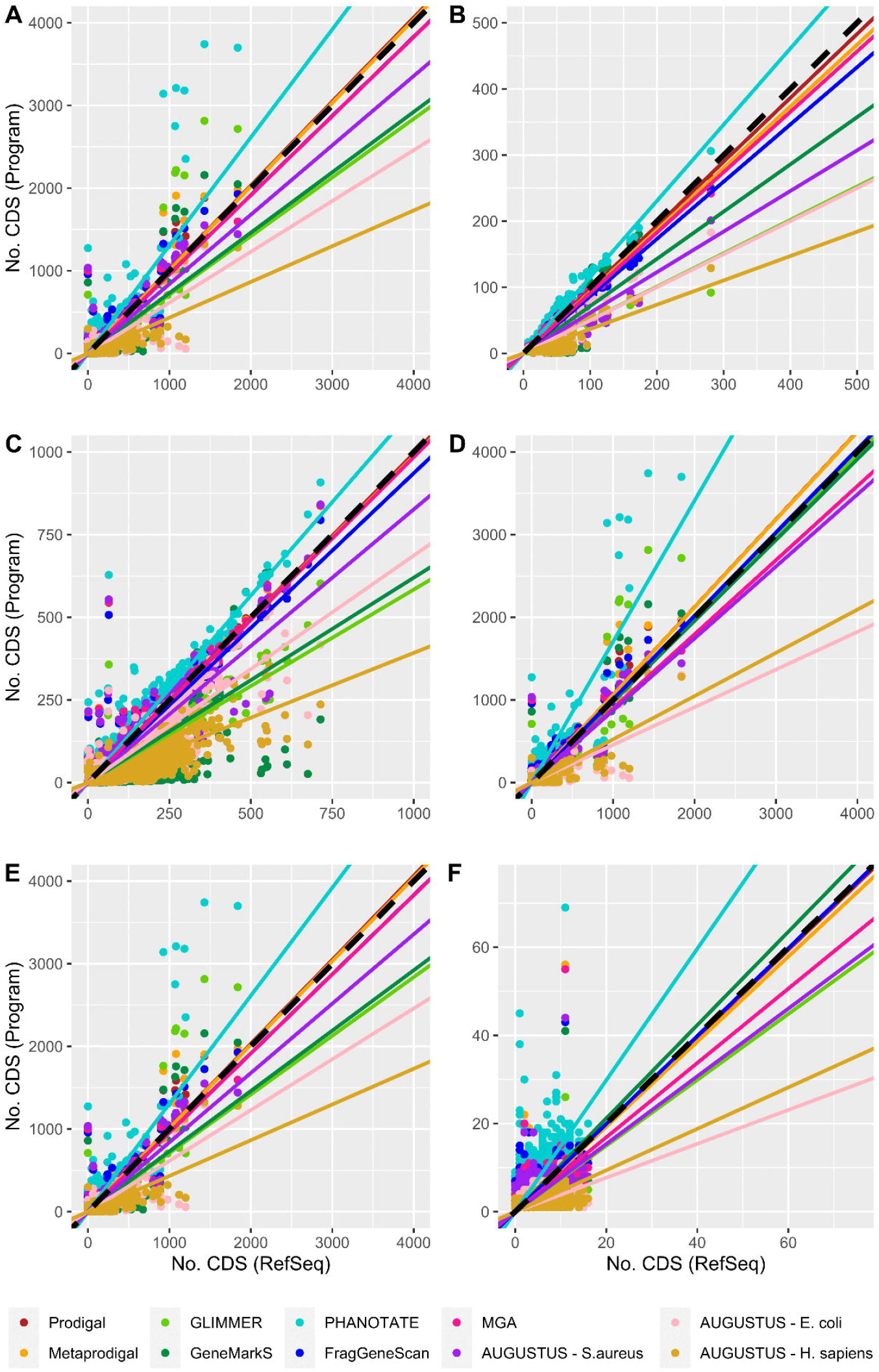
Correlation between the expected and observed number of coding sequences when considering (A) all known viral sequences, (B) viruses infecting archaea, (C) bacteriophages, (D) viruses infecting eukaryotes, (E) ds- and ss-DNA viruses, and (F) ds- and ss-RNA viruses. Dotted line is a 1:1 line.

### Coding Sequence Coordinate prediction

Secondly, to predict the quality of the coordinates of the viral coding sequences, F1 score, a measure that combines precision and sensitivity, was considered. Additionally, FDR and FNR were examined to evaluate the occurrence of false positives (i.e., false coordinates considered as true; type I error) and false negatives (i.e., true coordinates considered as false; type II error). Prodigal scored highly for all tests according to the F1 score (General: 83.26%; Viruses infecting Archaea: 80.02%; Bacteriophages: 86.25%; Viruses infecting Eukaryotes: 70.86%; ds- and ss-DNA viruses: 83.92%) except when analyzing RNA virus genomes (59.51%), where GeneMarkS obtained the best F1 score (60.84%), followed by Prodigal (59.51%) and Glimmer (56.60%). In contrast, for ds- and ss-DNA viruses, Prodigal (83.92%) generated the best results based on the F1 score, followed by Metaprodigal (81.91%) and MGA (80.60%). For both viruses infecting eukaryotes and bacteria, the highest FDR and FNR was associated with AUGUSTUS (median FDR [Viruses infecting eukaryotes]: 63.71 % ± 5.77 %; median FDR [Bacteriophages]: 29.40 % ± 33.14 %; median FNR [Viruses infecting eukaryotes]: 75.90 % ± 25.04 %; median FNR [Bacteriophages]: 44.77 % ± 0.50 %). Interestingly, the performance of the different ORF-calling programs to predict the quality of the coordinates in RNA virus genomes was very poor (median F1 score: 47.44% ± 46.92%; median precision: 45.05% ± 40.54%; median sensitivity: 52.46% ± 35.17%) compared to that in DNA viruses (median F1 score: 66.50% ± 43.59%; median precision: 75.19% ± 27.42%; median sensitivity: 63.69% ± 56.71%). In fact, GeneMarkS was more precise (64.26%) than other ORF-calling programs, including Prodigal (57.17%), for the prediction of the coordinates in RNA viruses. Overall, for all tests, the most sensitive ORF-calling program was Prodigal (Table 2).

**Table 2.** Accuracy, precision and sensitivity of the different programs. False Discovery Rate (FDR) and False Negative Ratio (FNR) are used to describe errors in the precision and sensitivity.

| Case | Program | F1 Score | Precision | Sensitivity | FDR<br>(Type I<br>Error) | FNR<br>(Type II<br>Error) |
| --- | --- | --- | --- | --- | --- | --- |
| All viruses (13,778) | PRODIGAL | 0.83261 | 0.826357 | 0.838958 | 0.1736427 | 0.1610417 |
|  | METAPRODIGAL | 0.810869 | 0.809406 | 0.812337 | 0.1905945 | 0.1876629 |
|  | GLIMMER | 0.374514 | 0.491625 | 0.302464 | 0.5083751 | 0.6975363 |
|  | GeneMarkS | 0.631786 | 0.747125 | 0.547296 | 0.2528753 | 0.4527039 |
|  | PHANOTATE | 0.689996 | 0.621607 | 0.775294 | 0.3783932 | 0.224706 |
|  | MGA | 0.793599 | 0.80448 | 0.783008 | 0.1955204 | 0.2169917 |
|  | FragGeneScan | 0.736218 | 0.750778 | 0.722212 | 0.2492221 | 0.2777881 |
|  | AUGUSTUS (*S. aureus*) | 0.564699 | 0.638428 | 0.506236 | 0.3615724 | 0.4937641 |
|  | AUGUSTUS (*E. coli*) | 0.585615 | 0.74484 | 0.482475 | 0.2551597 | 0.5175248 |
|  | AUGUSTUS (*H. sapiens*) | 0.226037 | 0.375398 | 0.161701 | 0.6246025 | 0.8382995 |
| Viruses infecting archaeal hosts (107) | PRODIGAL | 0.800237 | 0.810463 | 0.790266 | 0.1895369 | 0.2097341 |
|  | METAPRODIGAL | 0.794029 | 0.815076 | 0.774043 | 0.1849243 | 0.2259575 |
|  | GLIMMER | 0.357097 | 0.501361 | 0.277304 | 0.4986393 | 0.7226961 |
|  | GeneMarkS | 0.501389 | 0.693763 | 0.392541 | 0.3062371 | 0.6074594 |
|  | PHANOTATE | 0.709301 | 0.654914 | 0.773541 | 0.3450864 | 0.2264593 |
|  | MGA | 0.760603 | 0.793423 | 0.73039 | 0.206577 | 0.2696103 |
|  | FragGeneScan | 0.709419 | 0.759985 | 0.665161 | 0.2400153 | 0.3348386 |
|  | AUGUSTUS (*S. aureus*) | 0.494197 | 0.636676 | 0.403827 | 0.3633236 | 0.5961732 |
|  | AUGUSTUS (*E. coli*) | 0.38442 | 0.677612 | 0.268322 | 0.3223881 | 0.7316785 |
|  | AUGUSTUS (*H. sapiens*) | 0.183524 | 0.396226 | 0.119418 | 0.6037736 | 0.880582 |
| Bacteriophages (4,166) | PRODIGAL | 0.86248 | 0.862814 | 0.862146 | 0.1371862 | 0.1378536 |
|  | METAPRODIGAL | 0.854597 | 0.858627 | 0.850606 | 0.1413734 | 0.1493944 |
|  | GLIMMER | 0.347614 | 0.479301 | 0.272693 | 0.5206993 | 0.7273071 |
|  | GeneMarkS | 0.623633 | 0.760117 | 0.528702 | 0.2398835 | 0.471298 |
|  | PHANOTATE | 0.730571 | 0.683264 | 0.784917 | 0.3167362 | 0.2150832 |
|  | MGA | 0.851837 | 0.858461 | 0.845315 | 0.1415395 | 0.1546852 |
|  | FragGeneScan | 0.58717 | 0.563444 | 0.612983 | 0.4365558 | 0.3870173 |
|  | AUGUSTUS (*S. aureus*) | 0.61974 | 0.706003 | 0.552262 | 0.2939968 | 0.4477382 |
|  | AUGUSTUS (*E. coli*) | 0.652832 | 0.795405 | 0.553601 | 0.2045954 | 0.4463986 |
|  | AUGUSTUS (*H. sapiens*) | 0.204648 | 0.350757 | 0.144469 | 0.6492429 | 0.8555311 |
| Viruses infecting eukaryotic hosts (9,505) | PRODIGAL | 0.708596 | 0.679842 | 0.739891 | 0.3201583 | 0.2601094 |
|  | METAPRODIGAL | 0.626137 | 0.607279 | 0.646203 | 0.3927206 | 0.353797 |
|  | GLIMMER | 0.477917 | 0.528893 | 0.435904 | 0.4711074 | 0.5640963 |
|  | GeneMarkS | 0.670447 | 0.705276 | 0.638897 | 0.2947242 | 0.3611032 |
|  | PHANOTATE | 0.539783 | 0.427922 | 0.730823 | 0.572078 | 0.2691773 |
|  | MGA | 0.531444 | 0.552519 | 0.511917 | 0.4474812 | 0.4880828 |
|  | FragGeneScan | 0.58717 | 0.563444 | 0.612983 | 0.4365558 | 0.3870173 |
|  | AUGUSTUS (*S. aureus*) | 0.333522 | 0.36286 | 0.308572 | 0.6371399 | 0.6914276 |
|  | AUGUSTUS (*E. coli*) | 0.208491 | 0.347304 | 0.148956 | 0.6526962 | 0.8510445 |
|  | AUGUSTUS (*H. sapiens*) | 0.316537 | 0.460972 | 0.24102 | 0.5390285 | 0.7589803 |
| ds- and ss-DNA viruses (7,564) | PRODIGAL | 0.839228 | 0.833695 | 0.844835 | 0.1663046 | 0.1551654 |
|  | METAPRODIGAL | 0.819136 | 0.818105 | 0.82017 | 0.1818955 | 0.1798304 |
|  | GLIMMER | 0.3686 | 0.487136 | 0.296461 | 0.512864 | 0.7035386 |
|  | GeneMarkS | 0.632476 | 0.750597 | 0.546477 | 0.249403 | 0.4535226 |
|  | PHANOTATE | 0.697426 | 0.631266 | 0.779078 | 0.3687341 | 0.2209217 |
|  | MGA | 0.806025 | 0.817158 | 0.795191 | 0.1828416 | 0.2048086 |
|  | FragGeneScan | 0.742858 | 0.758995 | 0.727393 | 0.2410054 | 0.2726066 |
|  | AUGUSTUS (*S. aureus*) | 0.570832 | 0.64644 | 0.511059 | 0.3535598 | 0.4889415 |
|  | AUGUSTUS (*E. coli*) | 0.59314 | 0.753229 | 0.489172 | 0.2467711 | 0.5108278 |
|  | AUGUSTUS (*H. sapiens*) | 0.22311 | 0.373769 | 0.159014 | 0.626231 | 0.8409861 |
| ds- and ss-RNA viruses (6,214) | PRODIGAL | 0.595122 | 0.571708 | 0.620536 | 0.4282921 | 0.3794643 |
|  | METAPRODIGAL | 0.509863 | 0.499019 | 0.521189 | 0.5009811 | 0.4788114 |
|  | GLIMMER | 0.566089 | 0.610232 | 0.527901 | 0.3897682 | 0.4720989 |
|  | GeneMarkS | 0.608441 | 0.642596 | 0.577734 | 0.3574045 | 0.4222661 |
|  | PHANOTATE | 0.44324 | 0.345423 | 0.618343 | 0.6545772 | 0.3816568 |
|  | MGA | 0.333493 | 0.336816 | 0.330236 | 0.6631838 | 0.6697644 |
|  | FragGeneScan | 0.505519 | 0.483528 | 0.529606 | 0.5164718 | 0.4703945 |
|  | AUGUSTUS (*S. aureus*) | 0.341925 | 0.364488 | 0.321993 | 0.6355116 | 0.6780069 |
|  | AUGUSTUS (*E. coli*) | 0.121928 | 0.171686 | 0.094531 | 0.8283145 | 0.9054692 |
|  | AUGUSTUS (*H. sapiens*) | 0.324541 | 0.417475 | 0.265449 | 0.5825254 | 0.734551 |

## Discussion

In this study, we evaluated the performance of multiple ORF-calling programs for viral genome annotation based on the number of ORFs and their coordinates. According to our results, we found that viral gene predictions must be analyzed not considering the target host, but which nucleic acids the virus harbors. In fact, the differences in the performance of each program were more evident between ds- and ss-DNA viruses and ds- and ss-RNA viruses than among viruses infecting archaea, bacteriophages and viruses infecting eukaryotes.

We found that the performance of these ORF-calling programs was very poor for ds- and ss-RNA viruses, with GeneMarkS being the program that reached the highest F1 score, followed by Prodigal and Glimmer. This observation suggests the need for improvement for ORF calling programs to be able to deal with ds- and ss-RNA viruses, regardless of whether they are viruses infecting eukaryotes or prokaryotes. However, the vast majority of reported ds- and ss-RNA viruses infect eukaryotic organisms, driving the development of closed-reference homology-based bioinformatic tools, such as FLAN for influenza viruses (31), VIGOR for RNA viruses (32), ViPR and VAPID for human viruses (33,34), and VADR for non-flu viruses (35). Others have been developed for ss-DNA viruses, such as PuMA for papillomaviruses (36). The decision to develop and use a closed-reference homology-based method implies that the original viral references must be exceptionally well annotated. In this context, RNA and ss-DNA viruses harbor complex gene features with transcriptional and translational exceptions such as gene overlapping and alternative splicing, which are normally missed in most genome annotations (37,38). Additionally, from the perspective of bacteriophages, there is a considerable volume of ‘dark matter’ comprising poorly defined ORFs and genes of unknown function and there are very few examples of exceptionally well-annotated phage genomes (39). All these observations represent a major challenge for accurate and precise ORF-calling and gene annotation programs.

Considering the performance of the same programs applied to genome sequences from ds- and ss-DNA viruses, F1 scores were much higher than from RNA viruses. Prodigal reached the highest F1 score, followed by Metaprodigal and MGA. A potential explanation for this observation is the use of Prokka—a fast, de novo prokaryotic genome annotation pipeline—for the genome annotation of giant viruses, bacteriophages and viruses of Archaea, because this pipeline relies on Prodigal for the ORF calling process (9). Surprisingly, these results are not consistent with previously reported benchmarks, where MGA systematically generated less false positives than other ORF-calling programs (18) and GeneMarkS achieved the highest accuracy for the automatic gene identification for temperate phages due to the fewest number of false negatives and false positives (5). Nevertheless, no benchmarking has previously reported for the annotation of non-temperate lytic bacteriophage genomes, which are considered as an alternative to antibiotics to rapidly kill bacterial pathogens (“phage therapy”) (40). Additionally, it is important to note that none of the ORF-calling programs reached 90% accuracy for ds- and ss-DNA viruses, which is concordant with a previous benchmarking exercise (5). For this reason, several authors proposed the use of multiple ORF-calling programs to identify all viral genes (5,18,41). In such a way, it would be recommended to review the output of bioinformatic ORF prediction tools and manually interpret their findings (17,18,41), even though manual curation of an annotated genome is a time- and labor-intensive process. Of course, the ideal would be the manual curation of viral genomes, validated by wet-lab experiments to confirm the presence of these ORFs, as happens with RNA viruses, where the ORFs are characterized empirically via cDNA-gDNA hybridization (42-46) or using RNA-seq experiments (47-50). In the meantime, our evaluation of the current bioinformatic tools provides benchmarking to inform decisions about the most appropriate analysis pipelines for a given subject and highlights the need for more expression data to improve the rigor of reference genomes.

## Data availability

All Python 3 and R scripts used for this study are available at Github: https://github.com/EGTortuero/Benchmarking_ORF_calling_programs_in_viral_genomes

## Funding

This work was supported by the Biotechnology and Biological Sciences Research Council [grant numbers BB/T015616, BB/T016256].

## Conflict of Interest Disclosure

The authors declare that they have no competing interests.

## Acknowledgements

EGT wants to thank Beatriz Beamud (University of Valencia), Ramy Aziz (Cairo University) and Evelien Adriaenssens (Quadram Institute) for discussions on the benchmark design and interpretation.

## Notes

https://github.com/EGTortuero/Benchmarking_ORF_calling_programs_in_viral_genomes

### Competing Interest Statement

The authors have declared no competing interest.

## References

1. Ramamurthy, M., Sankar, S., Kannangai, R., Nandagopal, B. and Sridharan, G. (2017) Application of viromics: a new approach to the understanding of viral infections in humans. VirusDisease, 28, 349–359.

2. Miller, R.R., Montoya, V., Gardy, J.L., Patrick, D.M. and Tang, P. (2013) Metagenomics for pathogen detection in public health. Genome Medicine, 5, 81.

3. Simmonds, P., Adams, M.J., Benkő, M., Breitbart, M., Brister, J.R., Carstens, E.B., Davison, A.J., Delwart, E., Gorbalenya, A.E., Harrach, B. et al. (2017) Virus taxonomy in the age of metagenomics. Nature Reviews Microbiology, 15, 161–168.

4. Hayes, S., Mahony, J., Nauta, A. and Van Sinderen, D. (2017) Metagenomic Approaches to Assess Bacteriophages in Various Environmental Niches. Viruses, 9, 127.

5. Salisbury, A. and Tsourkas, P.K. (2019) A Method for Improving the Accuracy and Efficiency of Bacteriophage Genome Annotation. International Journal of Molecular Sciences, 20, 3391.

6. Roux, S., Adriaenssens, E.M., Dutilh, B.E., Koonin, E.V., Kropinski, A.M., Krupovic, M., Kuhn, J.H., Lavigne, R., Brister, J.R., Varsani, A. et al. (2019) Minimum Information about an Uncultivated Virus Genome (MIUViG). Nature Biotechnology, 37, 29–37.

7. Mitchell, A.L., Almeida, A., Beracochea, M., Boland, M., Burgin, J., Cochrane, G., Crusoe, M.R., Kale, V., Potter, S.C., Richardson, L.J. et al. (2019) MGnify: the microbiome analysis resource in 2020. Nucleic Acids Research.

8. Aziz, R.K., Bartels, D., Best, A.A., Dejongh, M., Disz, T., Edwards, R.A., Formsma, K., Gerdes, S., Glass, E.M., Kubal, M. et al. (2008) The RAST Server: Rapid Annotations using Subsystems Technology. BMC Genomics, 9, 75.

9. Seemann, T. (2014) Prokka: rapid prokaryotic genome annotation. Bioinformatics, 30, 2068–2069.

10. González-Tortuero, E., Sutton, T.D.S., Velayudhan, V., Shkoporov, A.N., Draper, L.A., Stockdale, S.R., Ross, R.P. and Hill, C. (2018), bioRxiv 277509.

11. Tisza, M.J., Belford, A.K., Domínguez-Huerta, G., Bolduc, B. and Buck, C.B. (2021) Cenote-Taker 2 democratizes virus discovery and sequence annotation. Virus Evolution, 7.

12. Delcher, A.L., Bratke, K.A., Powers, E.C. and Salzberg, S.L. (2007) Identifying bacterial genes and endosymbiont DNA with Glimmer. Bioinformatics, 23, 673–679.

13. Besemer, J. and Borodovsky, M. (1999) Heuristic approach to deriving models for gene finding. Nucleic Acids Research, 27, 3911–3920.

14. Besemer, J., Lomsadze, A. and Borodovsky, M. (2001) GeneMarkS: a self-training method for prediction of gene starts in microbial genomes. Implications for finding sequence motifs in regulatory regions. Nucleic Acids Research, 29, 2607–2618.

15. Zhu, W., Lomsadze, A. and Borodovsky, M. (2010) Ab initio gene identification in metagenomic sequences. Nucleic Acids Research, 38, e132–e132.

16. Hyatt, D., Chen, G.-L., Locascio, P.F., Land, M.L., Larimer, F.W. and Hauser, L.J. (2010) Prodigal: prokaryotic gene recognition and translation initiation site identification. BMC Bioinformatics, 11, 119.

17. Pope, W.H. and Jacobs-Sera, D. (2018) In Clokie, M. R. J., Kropinski, A. M.and Lavigne, R. (eds.), Bacteriophages: Methods and Protocols, Volume 3. Springer New York, New York, NY, pp. 217–229.

18. Lazeroff, M., Ryder, G., Harris, S.L. and Tsourkas, P.K. (2021) Phage Commander, an Application for Rapid Gene Identification in Bacteriophage Genomes Using Multiple Programs. PHAGE.

19. Daniel, W.W. (1995) Biostatistics : a foundation for analysis in the health sciences. 6th ed. ed. Wiley, New York ;.

20. Brister, J.R., Ako-Adjei, D., Bao, Y. and Blinkova, O. (2015) NCBI Viral Genomes Resource. Nucleic Acids Research, 43, D571–D577.

21. Tycowski, K.T., Guo, Y.E., Lee, N., Moss, W.N., Vallery, T.K., Xie, M. and Steitz, J.A. (2015) Viral noncoding RNAs: more surprises. Genes & Development, 29, 567–584.

22. McNair, K., Aziz, R.K., Pusch, G.D., Overbeek, R., Dutilh, B.E. and Edwards, R. (2018). Springer New York, pp. 231–238.

23. O’Leary, N.A., Wright, M.W., Brister, J.R., Ciufo, S., Haddad, D., McVeigh, R., Rajput, B., Robbertse, B., Smith-White, B., Ako-Adjei, D. et al. (2016) Reference sequence (RefSeq) database at NCBI: current status, taxonomic expansion, and functional annotation. Nucleic Acids Research, 44, D733–D745.

24. McNair, K., Zhou, C., Dinsdale, E.A., Souza, B. and Edwards, R.A. (2019) PHANOTATE: a novel approach to gene identification in phage genomes. Bioinformatics, 35, 4537–4542.

25. Hyatt, D., Locascio, P.F., Hauser, L.J. and Uberbacher, E.C. (2012) Gene and translation initiation site prediction in metagenomic sequences. Bioinformatics, 28, 2223–2230.

26. Rho, M., Tang, H. and Ye, Y. (2010) FragGeneScan: predicting genes in short and error-prone reads. Nucleic Acids Research, 38, e191–e191.

27. Noguchi, H., Taniguchi, T. and Itoh, T. (2008) MetaGeneAnnotator: Detecting Species-Specific Patterns of Ribosomal Binding Site for Precise Gene Prediction in Anonymous Prokaryotic and Phage Genomes. DNA Research, 15, 387–396.

28. Stanke, M., Diekhans, M., Baertsch, R. and Haussler, D. (2008) Using native and syntenically mapped cDNA alignments to improve de novo gene finding. Bioinformatics, 24, 637–644.

29. Brettin, T., Davis, J.J., Disz, T., Edwards, R.A., Gerdes, S., Olsen, G.J., Olson, R., Overbeek, R., Parrello, B., Pusch, G.D. et al. (2015) RASTtk: A modular and extensible implementation of the RAST algorithm for building custom annotation pipelines and annotating batches of genomes. Scientific Reports, 5, 8365.

30. R Core Team. (2021), R Foundation for Statistical Computing, Vienna, Austria.

31. Bao, Y., Bolotov, P., Dernovoy, D., Kiryutin, B. and Tatusova, T. (2007) FLAN: a web server for influenza virus genome annotation. Nucleic Acids Research, 35, W280–W284.

32. Wang, S., Sundaram, J.P. and Stockwell, T.B. (2012) VIGOR extended to annotate genomes for additional 12 different viruses. Nucleic Acids Research, 40, W186–W192.

33. Pickett, B.E., Sadat, E.L., Zhang, Y., Noronha, J.M., Squires, R.B., Hunt, V., Liu, M., Kumar, S., Zaremba, S., Gu, Z. et al. (2012) ViPR: an open bioinformatics database and analysis resource for virology research. Nucleic Acids Research, 40, D593–D598.

34. Shean, R.C., Makhsous, N., Stoddard, G.D., Lin, M.J. and Greninger, A.L. (2019) VAPiD: a lightweight cross-platform viral annotation pipeline and identification tool to facilitate virus genome submissions to NCBI GenBank. BMC Bioinformatics, 20.

35. Schäffer, A.A., Hatcher, E.L., Yankie, L., Shonkwiler, L., Brister, J.R., Karsch-Mizrachi, I. and Nawrocki, E.P. (2020) VADR: validation and annotation of virus sequence submissions to GenBank. BMC Bioinformatics, 21.

36. Pace, J., Youens-Clark, K., Freeman, C., Hurwitz, B. and Van Doorslaer, K. (2020) PuMA: A papillomavirus genome annotation tool. Virus Evolution, 6.

37. Chirico, N., Vianelli, A. and Belshaw, R. (2010) Why genes overlap in viruses. Proceedings of the Royal Society B: Biological Sciences, 277, 3809–3817.

38. Ashraf, U., Benoit-Pilven, C., Lacroix, V., Navratil, V. and Naffakh, N. (2019) Advances in Analyzing Virus-Induced Alterations of Host Cell Splicing. Trends in Microbiology, 27, 268–281.

39. Brum, J.R., Ignacio-Espinoza, J.C., Kim, E.-H., Trubl, G., Jones, R.M., Roux, S., Verberkmoes, N.C., Rich, V.I. and Sullivan, M.B. (2016) Illuminating structural proteins in viral “dark matter” with metaproteomics. Proceedings of the National Academy of Sciences, 113, 2436–2441.

40. Gordillo Altamirano, F.L. and Barr, J.J. Phage Therapy in the Postantibiotic Era. Clinical Microbiology Reviews, 32, e00066–00018.

41. Philipson, C., Voegtly, L., Lueder, M., Long, K., Rice, G., Frey, K., Biswas, B., Cer, R., Hamilton, T. and Bishop-Lilly, K. (2018) Characterizing Phage Genomes for Therapeutic Applications. Viruses, 10, 188.

42. Bornkamm, G.W., Desgranges, C. and Gissmann, L. (1983) In Bachmann, P. A. (ed.), New Developments in Diagnostic Virology. Springer Berlin Heidelberg, Berlin, Heidelberg, pp. 287–298.

43. Landry, M.L. and Fong, C.K. (1985) Nucleic acid hybridization in the diagnosis of viral infections. Clin Lab Med, 5, 513–529.

44. Hull, R. and Al-Hakim, A. (1988) Nucleic acid hybridization in plant virus diagnosis and characterization. Trends in Biotechnology, 6, 213–218.

45. Estes, M.K., Jiang, X., Zhou, Y.J. and Metcalf, T.G. (1990) In Bills, D. D., Kung, S.-D., Westhoff, D., Quebedeaux, B., Raleigh, E., Goss, J., Kotula, A.and Watada, A. (eds.), Biotechnology and Food Safety. Butterworth-Heinemann, pp. 185–191.

46. Younghusband, H.B., Egan, J.B. and Inman, R.B. (1975) Characterization of the DNA from bacteriophage P2-186 hybrids and physical mapping of the 186 chromosome. Molecular and General Genetics MGG, 140, 101–110.

47. Bernal-Vicente, A., Donaire, L., Torre, C., Gómez-Aix, C., Sánchez-Pina, M.A., Juarez, M., Hernando, Y. and Aranda, M.A. (2018) Small RNA-Seq to Characterize Viruses Responsible of Lettuce Big Vein Disease in Spain. Frontiers in Microbiology, 9.

48. Liu, C., Liu, Y., Liang, L., Cui, S. and Zhang, Y. (2019) RNA-Seq based transcriptome analysis during bovine viral diarrhoea virus (BVDV) infection. BMC Genomics, 20, 774.

49. Wicke, L., Ponath, F., Coppens, L., Gerovac, M., Lavigne, R. and Vogel, J. (2021) Introducing differential RNA-seq mapping to track the early infection phase for Pseudomonas phage LKZ. RNA Biology, 18, 1099–1110.

50. Li, T., Zhang, Y., Dong, K., Kuo, C.-J., Li, C., Zhu, Y.-Q., Qin, J., Li, Q.-T., Chang, Y.-F., Guo, X. et al. (2020) Isolation and Characterization of the Novel Phage JD032 and Global Transcriptomic Response during JD032 Infection of Clostridioides difficile Ribotype 078. mSystems, 5, e00017–00020.

